# Identification of Transcription Factor Binding Sites using ATAC-seq

**DOI:** 10.1101/362863

**Authors:** Zhijian Li, Marcel H. Schulz, Martin Zenke, Ivan G. Costa

**Affiliations:** Institute for Computational Genomics, RWTH Aachen University Medical School, Aachen, Germany.; Department of Cell Biology, Institute of Biomedical Engineering, RWTH Aachen University Medical School, Aachen, Germany.; Cluster of Excellence for Multimodal Computing and Interaction, Saarland Informatics Campus, Saarland University, Saarbrücken, Germany; Computational Biology & Applied Algorithmics, Max Planck Institute for Informatics, Saarbruücken, Germany; Helmholtz Institute for Biomedical Engineering, RWTH Aachen University, Aachen, Germany; Aachen Institute for Advanced Study in Computational Engineering Science (AICES), RWTH Aachen University, Aachen, Germany

**Keywords:** computational footprinting, open chromatin, ATAC-seq, cleavage bias, strand-specific bias

## Abstract

Transposase-Accessible Chromatin (ATAC) followed by sequencing (ATAC-seq) is a simple and fast protocol for detection of open chromatin. However, computational footprinting in ATAC-seq, i.e. search for regions with depletion of cleavage events due to transcription factor binding sites, has been poorly explored so far. We propose HINT-ATAC, a footprinting method that addresses ATAC-seq specific protocol artifacts. HINT-ATAC uses a probabilistic framework based on Variable-order Markov models to learn the complex sequence cleavage preferences of the transposase enzyme. Moreover, we observed specific strand specific cleavage patterns around the binding sites of transcription factors, which are determined by local nucleosome architecture. HINT-ATAC explores local nucleosome architecture to significantly outperform competing footprinting methods in predicting transcription factor binding sites by ChIP-seq. HINT-ATAC is an open source software and available online at www.regulatory-genomics.org/hint

## 2 Introduction

DNase I hypersensitive sites sequencing (DNase-seq) has become the standard choice for genome-wide detection of open chromatin (Boyle et al. 2008; Crawford et al. 2006b; Neph et al. 2012a; Vierstra and Stamatoyannopoulos 2016). In this protocol, the start of DNA fragments reflects the exact location of DNase I cleavage, therefore transcription factor binding sites (TFBS) leave footprints (depletion of DNase-seq signals) and these indicate the location of TFBS. Assays for Transposase-Accessible Chromatin followed by sequencing (ATAC-seq) have been proposed as an alternative to DNase-seq (Buenrostro et al. 2013). ATAC-seq requires fewer cells (down to 500) and is less laborious than DNase-seq (Buenrostro et al. 2013). A recent variant of the protocol (Omni-ATAC) increases the proportion of reads from non-mithocondrial DNA and allows working of frozen tissues (Corces et al. 2017). A similar number of studies based on ATAC-seq and DNase-seq (247 and 210 respectively) have been deposited in Gene Expression Omnibus in the last year, while there are three times more ATAC-seq samples than DNase-seq samples in these studies^1^. This indicates the emerging importance of ATAC-seq and confirms that its experimental simplicity makes it a choice for studies with large number of samples as in clinical scenarios (Rendeiro et al. 2016).

Computational footprinting methods are crucial in the analysis of DNase-seq (Vierstra and Stamatoyannopoulos 2016; Gusmao et al. 2016). Among others, footprinting on DNase-seq data has been used to detect regulatory lexicon of several cells (Neph et al. 2012b; Schmidt et al. 2017; Ho et al. 2017; Kolovos et al. 2016; Lin et al. 2015), investigate responses to environmental stimuli (Goldstein et al. 2017) and detection of variants impacting transcription factor binding (Schwessinger et al. 2017). While footprints are also observed in ATAC-seq data (Buenrostro et al. 2013), computational footprinting of ATAC-seq data has been poorly addressed for far. The few studies contrasting ATAC-seq and DNase-seq show that ATAC-seq footprints have a slightly inferior accuracy than DNase-seq based footprints (Quach and Furey 2016) or leave more unclear footprint profiles than DNase-seq (Schwessinger et al. 2017). Moreover, most work performing footprinting in ATAC-seq so far (Buenrostro et al. 2013; Raj et al. 2015; Quach and Furey 2016) treated ATAC-seq and DNase-seq similarly and ignored characteristics intrinsic to the ATAC-seq protocol.

A possible reason for the lower performance of ATAC-seq is its cleavage enzyme Tn5, which has a large DNA binding site and a more complex cleavage mechanism than DNase-I (Fig. 1A & B). The Tn5 dimer cleaves the DNA by inserting two distinct adapters in the DNA fragment ends. Cleavage leaves two 9bps single strand DNA ends that are later extended in the ATAC-seq protocol (Fig. 1C). Tn5 displays a large palindromic (17bp) “transposase motif” (Reznikoff 2008; Buenrostro et al. 2013) in the start of aligned ATAC-Seq reads (Fig. 1A). The palindromicity reflects the fact that Tn5 works in dimmers and individual proteins bind to DNA in reverted orientations. Moreover, the motif is centered around position +5, which indicates Tn5 binds to the middle of the 9bp single stranded DNA. In contrast, DNase-I leaves a short motif close to the start of reads in DNase-seq experiments (Fig. 1B).

**Figure 1:**
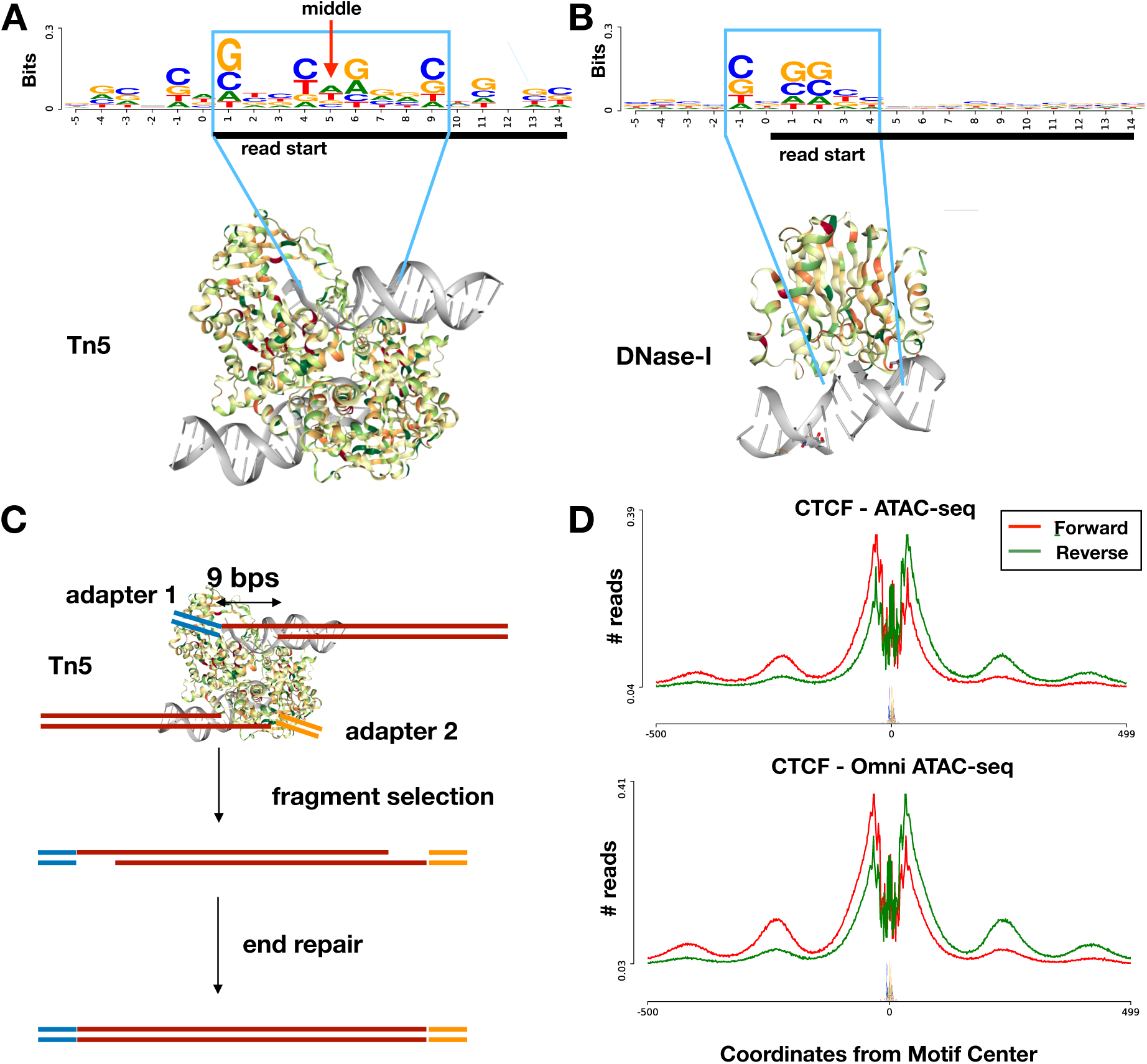
Sequence motif relative to aligned read start after cleavage with Tn5 and DNase I enzymes on naked DNA ATAC-seq (A) and DNase-seq (B) experiments. The size of the motifs are reflected by the structural protein contacts of Tn5 and DNase-I (Protein Data Bank entries 1MM8 and 2DNJ). C - Tn5 inserts adapters in both DNA ends. Moreover, DNA is cleaved in two 9 bps single ends, which are later repaired in the ATAC-seq protocol. D - Cleavage profiles around CTCF Chip-seq peaks indicate strand specific cleavage preference left/right of the TF binding site for both standard and Omni ATAC-seq libraries. Smaller peaks far from the TF binding represent linker regions between histones

The DNA binding preferences of enzymes cause the so called sequence specific cleavage bias. Computational bias correction is an important aspect of the analysis of DNase-seq and ATAC-seq data (He et al. 2014; Sung et al. 2014; Martins et al. 2017). All bias correction methods proposed so far are based on bias estimates of *k*-mer sequences around the start of aligned reads, i.e., probability of finding a 6-mer around the start of DNase-seq reads vs. the probability of the k-mer in the genome (He et al. 2014; Sung et al. 2014; Gusmao et al. 2016; Wang et al. 2017; Schwessinger et al. 2017). These methods require the estimation of a multinomial distribution and will suffer from over-fitting for large *k* as possibly required for ATAC-seq data. A solution for dealing with large k-mer size is the use of position weight matrices (PWMs) which is the most popular model for protein-DNA interaction (Stormo et al. 1982) and assumes statistical independence between positions. Variable order Markov models (VOM) represent an alternative to PWMs, where relevant dependencies are learned from the data (Ben-Gal et al. 2005; Keilwagen and Grau 2015). Currently, we are unaware of any work applying VOMs for modeling the bias of ATAC-seq or any cleavage enzymes.

The large and asymmetric nature of the Tn5 dimmer (Fig 1A) will also make cleavage dependent on structural features of the neighboring proteins (transcription factors or histones) and size of accessible DNA (Schep et al. 2015). Fragment size distribution of ATAC-seq libraries indicates indeed that cleavage between the small linker DNA between nucleosomes is possible, but less likely than cleavage of short fragments from open chromatin (Buenrostro et al. 2013). Indeed, strand specific ATAC-seq profiles around CTCF, which is a TF usually binding alone and surrounded by equally spaced histones (Buenrostro et al. 2013; Vierstra et al. 2014), has an intricate cleavage pattern with higher number of cleavage events in the forward strand left of the TF binding site for both major ATAC-seq protocols (Fig. 1D). The strand specificity is particularly high in linker regions. While strand specificity has been previously observed in DNase-seq (Piper et al. 2013; Schwessinger et al. 2017), the strand specific nature of ATAC-seq and the influence of local nucleosome architecture has not been shown or explored until date.

We propose here HINT-ATAC, the first footprinting method dealing with ATAC-seq specific characteristics. First, HINT-ATAC includes a probabilistic approach for correction of cleavage bias based on a specific type of Variable order Markov models (Keilwagen and Grau 2015). We perform a comprehensive evaluation of distinct bias correction approaches (k-mer based, PWM and VOM) for ATAC-seq and DNase-seq data under distinct word sizes. Bias correction strategies are evaluated in the ability to recover TF ChIP-seq binding sites as in Cuellar-Partida et al. (2011); Gusmao et al. (2016). Next, we explore the fact that fragment sizes, as indicated by paired-end reads, allow the decomposition of cleavage signals by fragments potentially containing zero, one or more nucleosomes (Vierstra et al. 2014; Buenrostro et al. 2013). We used distinct combinations of strand specific and nucleosome decomposed signals to generate multivariate cleavage profiles used as input to the Hidden Markov model. We use data of independent cell lines to contrast the performance of HINT-ATAC with state of art footprint methods DNase2TF (Sung et al. 2014), PIQ (Sherwood et al. 2014), HINT (Gusmao et al. 2014) and DeFCoM (Quach and Furey 2016). HINT-BC has been previously proposed by us for footprinting in DNase-seq data, while DeFCoM is the only footprint method evaluated in ATAC-seq up to date. We also contrast the performance of HINT-ATAC in distinct variants of ATAC-seq and DNase-seq protocols.

## 3 Results

### 3.1 Variable order Markov models improve cleavage bias correction

We investigate here the impact of distinct methods for bias corrections (k-mer, PWM and VOM) to improve footprinting in ATAC-seq and also in DNase-seq protocols. These steps are performed previous to the application of HINT (Gusmao et al. 2014) (see Methods Section). Besides the standard approach to use reads from the ATAC-seq/DNA-seq library itself, we also evaluate the use of bias estimates from ATAC-seq/DNase-seq libraries performed in the deproteinated DNA (naked DNA). Methods were evaluated on their recovery of footprints with binding sites supported by ChIP-seq peaks following Gusmao et al. (2016). We use here ChIP-seq of 32 TF on GM12878 cells (**`train dataset`**). For each TF, we calculate the area under precision recall curve (AUPR) and under the reciever operating characteristics curve (AUC) for distinct false positive rates (1%, 10% and 100%) A final ranking score is obtained by combining the ranking of a method for each of the six statistics. A higher ranking score indicates higher recovery of ChIP-seq supported footprints than compared methods.

A first question is the influence of the word size (*k*) within each bias correction method. For DNase-seq data, bias correction methods performed best with small word size (4-8), while larger words were best for ATAC-seq (8-12). These observations fits with the size of motifs detected for DNase-I and Tn5 cleavage sites (Fig. 1A & B). Next, we compared distinct bias correction methods, where k was fixed to the best choices for each method (Fig. 2A & B). VOM models with *k* = 8 were ranked best in both ATAC-seq and DNase-seq protocols. It significantly outperformed PWM, K-mer (naked DNA) and uncorrected signal for ATAC-seq (*p*-value < 0.05; Friedman-Nemenyi test) and the uncorrected signal for DNase-seq (*p*-value < 10^−7^; Friedman-Nemenyi test). We also observed that estimates based on naked DNA obtained lower ranks than their counterparts based on reads from the ATAC/DNase-seq library at hand. Therefore, we adopt the use of VOM(8) bias correction as standard in HINT-ATAC and in all experiments below.

**Figure 2:**
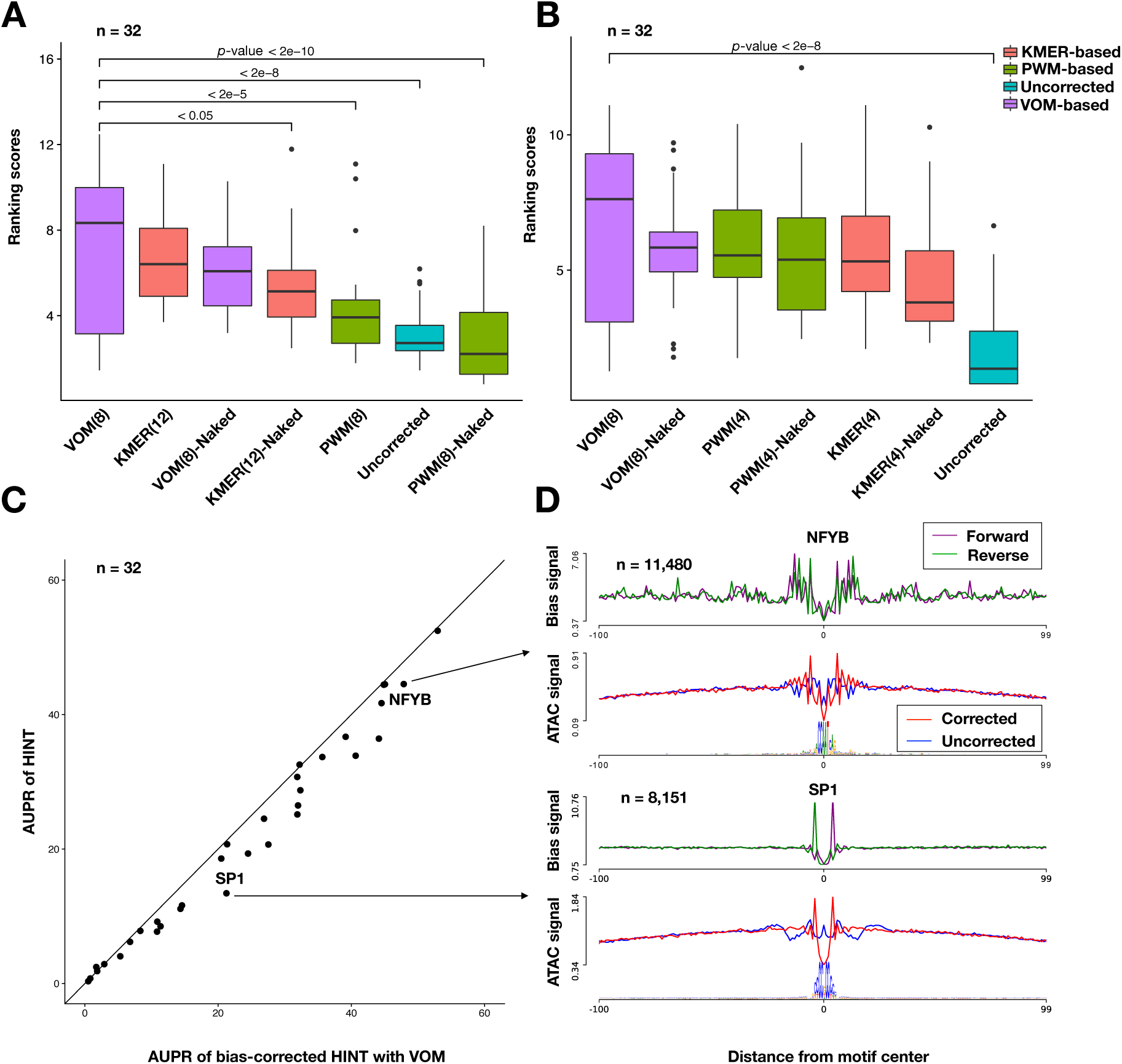
Comparison of bias estimation methods in ATAC-seq (A) and DNase-seq (B) on 32 TF ChIP-seq data from GM12878. *Y*-label indicate the ranking score, where higher values indicates higher recovery of footprints supported by TF ChIP-seq peaks. (C) The scatter plot contrasting AUPR of HINT with VOM-based estimation with 8-mer (*x*-axis) and HINT without bias correction (*y*-axis). NFYB and Sp1, which are TFs among the ones with highest AUPR increase, have a clear improvement of footprint profiles after correction (D).

The AUPR of HINT with VOM(8) bias correction is higher than the AUPR of HINT with uncorrected signal for 29 of 32 TFs investigated, as exemplified by the clear footprint profiles of NFYB and Sp1 after bias correction (Fig. 2C). AUPR only decreased marginally (average of 0.003) for 3 TFs. Lastly, we evaluated the similarity between sequence bias estimates (based on VOM 8-mer approach) on 31 ATAC-seq libraries. We observed that all ATAC-seq libraries based on the standard ATAC and Omni-ATAC protocol group together and have quite similar bias (average R=0.96). Bias estimates based on naked DNA also grouped with standard ATAC-seq libraries, but showed a smaller average correlation (R=0.88) with other standard libraries. Single cell ATAC-seq libraries grouped apart for all other experiments (average R=0.82 in relation to standard libraries). The distinct bias in protocols with few cells might be an effect of over-digestion of chromatin by Tn5 (Buenrostro et al. 2015). Given that cleavage bias varies in distinct degrees for each library, these results support the use of bias estimates based on reads from the ATAC-seq library at hand.

### 3.2 Strand specific and nucleosome decomposition improves footprint predictions in ATAC-seq

We evaluate here variants of HINT-ATAC HMM models in their abilities to predict footprints on the standard and Omni ATAC protocols. Of particular interest is to explore the use of strand specific signals and local nucleosome architecture for footprint prediction. We decompose cleavage signals by only considering reads from nucleosome free fragments or reads from fragments with particular number of nucleosomes. Decomposition is based on fragment size intervals defined on the distribution of paired-end aligned reads (Fig. 3A). Here, we evaluate the following nucleosome decompositions as input for HINT-ATAC: all reads (**`all`**); only nucleosome free reads (**`Nfr`**); joint analysis of nucleosome free and nucleosome containing signals (**`Nfr`** & **`+1N`**); and the joint analysis of nucleosome free, one nucleosome and two or more nu-cleosomes signals (**`Nfr`** & **`1N`** & **`+2N`**). For each decomposition, we have two signals (number of reads and slope of number of reads (Gusmao et al. 2014)) for both positive and negative strands. Therefore, strand-specific nucleosome decomposition strategies produce signals with 4 (**`All and Nfr`**), 8 (**`Nfr`** & **`+1N`**) and 12 dimensions (**`Nfr`** & **`1N`** & **`+2N`**). We then evaluate models for standard ATAC-seq and Omni-ATAC-seq separately on the **`training dataset`** with 32 TFs in GM12979 cells.

**Figure 3:**
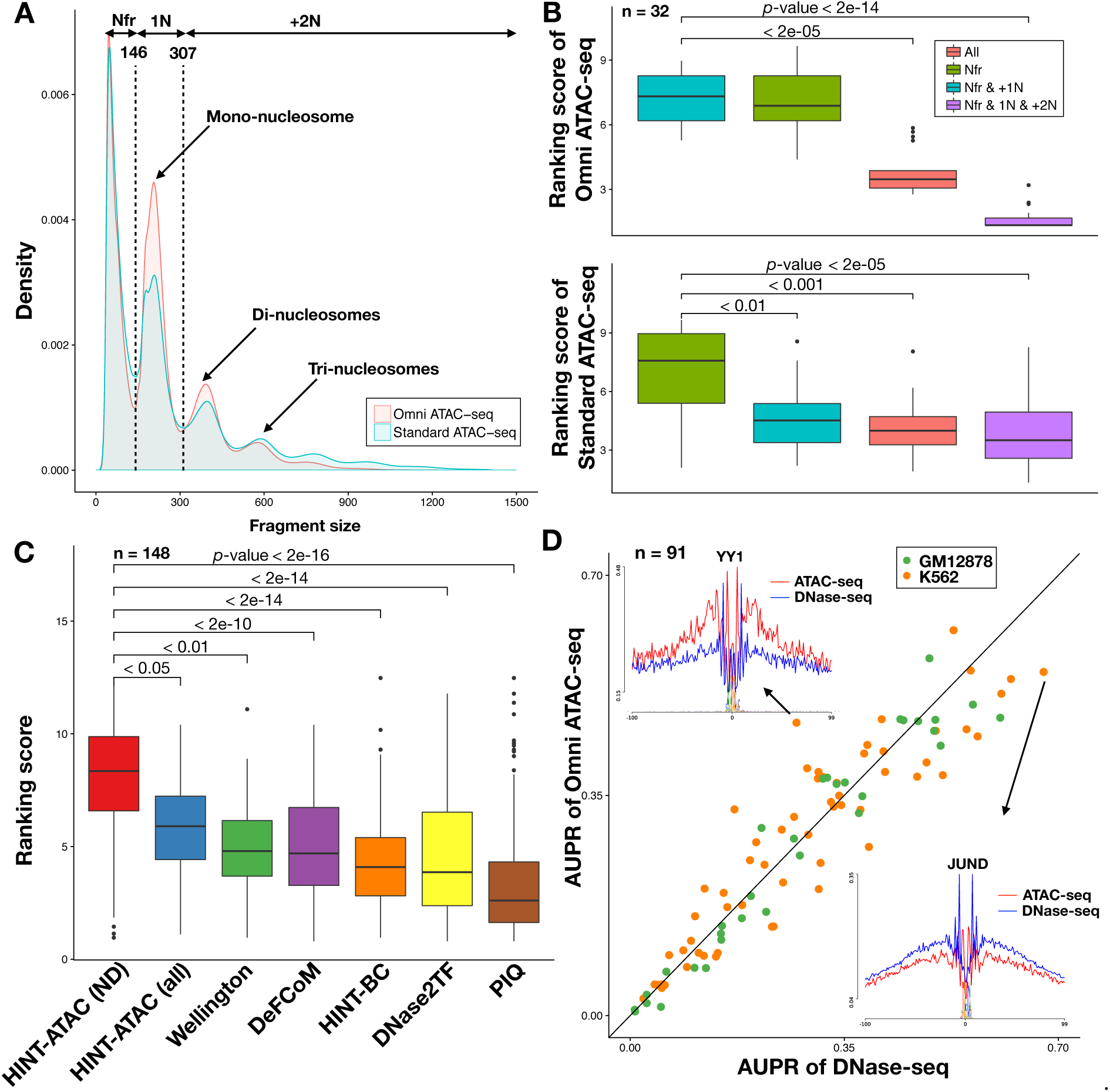
A - Fragment size distribution for Omni and standard ATAC-seq on GM12878 cells indicate clear peaks representing fragments with particular numbers of nucleosomes. Local minimum values were used to define nucleosome free fragments **`Nfr`**, fragments with one nucleosome **`1N`** and fragments with one or more **`+2N`** nucleosomes. B - Comparison of HINT-ATAC models with distinct nucleosome decomposition strategies of Omni ATAC-seq (top) and standard ATAC-seq (bottom) on **`training dataset`** (GM12787 cell). Higher ranking score indicates highest recovery of ChIP-seq supported binding sites. C - Comparative evaluation of HINT-ATAC with **`all`** reads and with nucleosome decomposition (ND), HINT, Wellington, DNase2TF, DeFCoM and PIQ in the **`test dataset`** (H1-ESC and K562 cells). D - AUPR values of DNase-seq (double hit protocol) vs ATAC-seq (Omni) for 91 factors, of which 41 factors obtain higher AUPR using ATAC-seq. The footprint profiles of two factors with the highest AUPR difference are shown.

For each nucleosome decomposition strategy (**`All`**, **`Nfr`**, **`Nfr`** & **`1N`**,**`Nfr`** & **`1N`** & **`+2N`**) we evaluated the optimal number of states and the use of strand-specific signal on the **`training dataset`**. While the optimal number of states was specific to the decomposition strategy, models with strand-specific signals had highest ranking in 7 out of 8 comparisons. Next, we contrasted the performance of the best models for each decomposition (Fig. 3B). The use of nucleosome free reads (**`Nfr`**) obtained higher ranking scores than other models for standard ATAC, while the use of **`Nfr`** & **`+1Nr`** or only **`Nfr`** signals obtained higher ranking scores than other models for Omni-ATAC (*p*-value < 0.01; Friedman-Nemenyi test). This indicates that separating cleavage signals by the presence and number of nucleosomes between two cleavage events is better than using all reads for finding footprints in ATAC-seq. The HMM with 9 states and strand specific **`Nfr`** reads will be adopted as the HINT-ATAC model for standard ATAC and the HMM with 7 states based on **`Nfr`** & **`+1N`** strand specific signals will be adopted for Omni-ATAC protocol.

We used the **`test dataset`** based on K562 and H1-ESC cells for standard ATAC-seq and K562 for Omni-ATAC (148 TFs in total) to compare the performance of HINT-ATAC with state-of-the-art footprinting methods. This includes DNase2TF, PIQ, Wellington and HINT-BC, which have been reported to perform well with DNase-seq data (Gusmao et al. 2016) and DeFCoM, which has been recently evaluated in ATAC-seq data. We also included the evaluation of a version of HINT-ATAC with **`all`** reads, as nucleosome decomposition is not possible for ATAC-seq libraries sequenced with single ends. Both versions of HINT-ATAC had the highest ranking scores (Fig. 3C) and significantly outperformed all other methods (*p*-value < 0.01; Friedman-Nemenyi test). Similar rankings were observed when only considering Omni-ATAC and standard ATAC libraries independently. HINT-ATAC based on nucleosome decomposition significantly outperformed the use of **`all`** reads (*p*-value < 0.05; Friedman-Nemenyi test). Interestingly, the only other method using strand specific information—Wellington— was ranked third and significantly outperformed its lower-ranked competitors (*p*-value < 0.01; Friedman-Nemenyi test).

### 3.3 Footprints based on Omni-ATAC-seq and DNase-seq have similar predictive performance

Next, we compared the performance of distinct ATAC-seq protocols (standard (Buenrostro et al. 2013), fast (Corces et al. 2016) and Omni (Corces et al. 2017) ATAC-seq) and cell number (bulk 50.000, 500 or single cells) (Buenrostro et al. 2015) in either GM12878, K562 and H1-ESC cell lines. To contrast results with DNase-seq based footprints, we have also optimized HINT models to consider strand specific signals for DNase-seq of single hit (Sabo et al. 2004) and double hit (Crawford et al. 2006a) protocols. Note that nucleosome decomposition is not possible here due to lack of paired end libraries. We observed that footprint prediction, on either double or single hit DNase-seq, benefits from the use of strand specific signals (*p*-value < 0.05;; Friedman-Nemenyi test). Therefore, we adopted strand specific signals and an HMM with 9 states for double hit DNase-seq and 7 states for single hit DNase-seq from here on. These models, which also include VOM based bias correction, will be refereed as HINT-DNase.

Double hit DNase-seq and Omni ATAC-seq have significantly higher ranking scores than other protocols (*p*-value < 0.05; Friedman-Nemenyi). Overall, standard ATAC-seq and fast-ATAC-seq protocols had lowest ranking scores. Down-sampling of libraries with a high number of reads did not influence rankings. On an individual factor level, we observed that TFs with higher AUPR on ATAC-seq have more ATAC-seq cleavage sites surrounding the binding site and vice-versa, as exemplified by YY1 and JunD for both Omni-ATAC (Fig. 3D) or standard ATAC protocols. We also investigated if differences between ATAC-seq and DNase-seq might arise from the previously reported lower performance of Tn5 in digesting enhancer regions (Sos et al. 2016; Corces et al. 2017). We therefore split our dataset with histone based annotation of promoter and enhancer regions from chromHMM (Ernst and Kellis 2012). We observed that the AUPR difference (DNase-seq - ATAC-seq) is significantly larger for enhancer regions than in promoter regions (*p*-value < 0.01; Wilcoxon signed rank test) with standard ATAC-seq, however no such difference was found in Omni-ATAC-seq. DNase-seq obtained significantly higher AUPR for all 21 TFs from the bZIP family for both ATAC protocols and for factors of the helix-loop-helix family for standard ATAC (*p*-value < 0.001; Mann-Whitney test). This is an indication that structural features of the TF, which are usually shared by TF families, negatively affect the ability of Tn5 to cleave DNA around specific TFs.

### 3.4 Local nucleosomes architecture and strand specific ATAC-seq digestion profiles

To understand the importance of nucleosome number decomposition and strand specific bias in footprint prediction, we analyzed the average cleavage profiles of distinct decomposed signals around CTCF ChIP-seq peaks in GM12878 cells (Fig. 4A and 4B). First, we calculated the position of linkers (by finding peak centers of **`1N`** and **`+2N`** reads) and then the middle position of histones. The intervals between histone and CTCF center were used to count the number of reads associated to linkers and on the open chromatin region left or right of CTCF. Of particular interest are cleavage events left/right to CTCF, as these are the ones delineating footprints. While profiles based on all reads are strand specific in the vicinity of CTCF (1.38 ratio of forward/reverse reads left of CTCF binding), we observed a stronger strand cleavage bias in nucleosome free fragments in both Omni-ATAC-seq (2.6 ratio) and standard ATAC-seq (2.3 ratio). As shown in Fig. 4C, nucleosome free reads can arise from DNA fragments with (type I) or without (type II) CTCF binding in the middle. As sequencing is performed in the 5’ to 3’ direction, type I reads will generate forward reads left to CTCF, while type II reads will generate both forward and reverse reads left to CTCF. Using CTCF binding sites supported by ChIP-seq to count type I and II **`Nfr`** fragments, we detected more type I than type II reads in **`Nfr`** fragments for Omni-ATAC or standard ATAC, which supports the bias towards forward reads in the regions adjacent to the left of CTCF binding and vice-versa.

**Figure 4:**
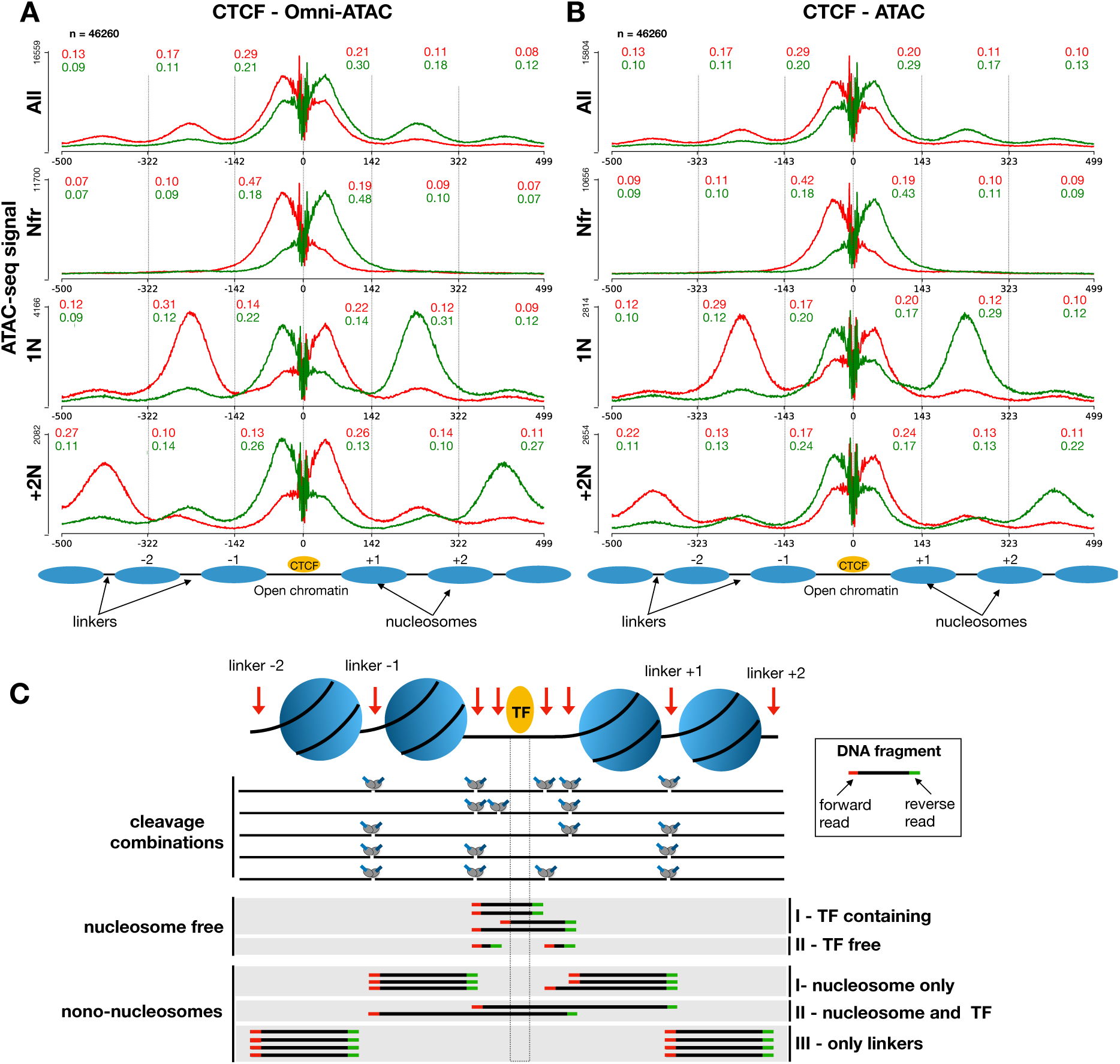
The average bias corrected cleavage profile for distinct nucleosome number decompositions around CTCF ChIP-seq peaks for Omni-ATAC (A) and standard ATAC (B) protocols in GM12878 cells. To quantify the amount of strand bias relative to CTCF, we estimate the proportion of reads in forward (red) and reverse (green) around intervals corresponding to the center of nucleosomes and CTCF (positions −322, −142, 0, 142, 322). Estimates of strand bias are the ratio of forward and reverse reads in a particular region, i.e. ratio of 0.29/0.21=1.38 for the internal [-142,0] for **`All`** signals for Omni-ATAC-seq. C - Tn5 digests open chromatin region left/right to the TF binding or nucleosome linkers. Nucleosome free reads will generate fragments with (type I) or without (type II) the TF bound to DNA. As sequencing is performed from the 5’ to 3’ ends, fragments will always generate forward reads in the left (red) and reverse reads in the right (green) of the fragments. DNA fragments from **`1N`** decomposition contain either only a histone (types I and III) or a histone and a TF (type II). Type I fragments will produce reads of the reverse strand left to the TF (forward strand right to the TF), while type II forward reads left to the TF. Type I, II and III reads can be defined analogously to fragments with two or more nucleosomes.

While **`Nfr`** reads have few cleavage events on linker regions, we observed clear peaks on linkers −1 and + 1 for **`1N`** reads and clear peaks on linkers −2 and +2 for **`+2N`** reads (Fig. 4A and B). This confirms the soundness of the nucleosome number decomposition strategy in separating reads regarding the potential presence of distinct numbers of nucleosomes using only fragment sizes. Interestingly, the strand bias in the region left to CTCF is reverted for **`1N`** (ratio 0.63) and **`+2N`** signals (ratio 0.5) in contrast to **`Nfr`** reads, while cleavage events in the linker −1 (for **`1N`** reads) and linker −2 (for **`+2N`** reads) are biased towards forward strand (ratio 1.9-2.6). As proposed in Fig. 3C, **`1N`** (and **`+2N`** reads) can be divided into three types: fragments including one nucleosome close to CTCF (type I), fragments containing a nucleosome and CTCF (type II) or others (type III). We observed more type I reads in comparison to type II (or type III) in Omni-ATAC-seq and standard ATAC-seq for both **`1N`** and **`+2N`**. The higher number of type I reads, which generates reverse strand reads left of CTCF, explains the inverted strand cleavage bias observed in **`1N`** and **`+2N`** reads (Fig. 4C).

CTCF is an exceptional TF, which usually binds alone to the DNA and has a tight distance to the next nucleosome (Buenrostro et al. 2013; Vierstra et al. 2014). While peaks associated to cleavage in linkers regions are less prominent in other TFs and the distance to the next linker varies for each TF (Buenrostro et al. 2013), we observed the similar strand bias for nucleosome number decomposed signals for other TFs with or without cleavage bias correction. Finally, we investigated if TF specific properties, as the distance to the next linker, the amount of strand bias on average profiles is related to the AUPR values of HINT-ATAC. While there is no clear association between distance to nucleosomes and AUPR, we observe that the AUPR of TF binding in enhancers is correlated with strand specificity for all ATAC and DNase-seq protocols (average R=0.57). Altogether, these results indicate that properties of the cleavage events generating fragments (nucleosome free, nucleosome containing with or without the TF) introduce a strand specific cleavage bias relative to the TF binding site. These strand bias are crucial in the delineation of footprints.

## 4 Discussions

We present here the first computational footprinting method tailored for the ATAC-seq protocol. First, we show that the use of Variable-order Markov models (VOM) for correction of cleavage bias is crucial for ATAC-seq. This approach, which estimates dependencies between sequence positions from the data, is able to model cleavage bias spanning larger motifs more accurately than the k-mer based approach because it avoids over-fitting. Previously, Schwessinger et al. (2017) reported difficulty in detection of footprints around motifs associated to regulatory variants in ATAC-seq data. As a practical example of the power of VOM models, we observe that bias correction improves footprint profiles for the majority of these motifs. There is a growing attention in the field by the fact that cleavage bias is present in any sequencing protocol using cleavage/digestion enzymes (Madrigal 2015; Foulk et al. 2015; Wang et al. 2017). For example, the “digestion bias” of nucleases has been shown to induce artifacts in nascent RNA-seq protocols (Foulk et al. 2015). The same enzyme is used in ChIP-seq variants (ChIP-exo (Rhee and Pugh 2011); ChIP-nexus (He et al. 2015)) and is likely to influence the detection of ChIP-exo footprints. VOM models represent a flexible framework for modeling cleavage bias, which is likely to improve downstream analysis of any of these protocols.

The strand specific digestion patterns around transcription factor binding sites represent another overseen aspect of ATAC-seq. Decomposition of DNA fragments by nucleosome number contains intricate strand specific cleavage patterns relative to TF binding. We show that the strand bias of ATAC-seq protocols arises from a preference of having fragments with the TF alone or fragments containing only nucleosomes with a digestion event close to the TF (Fig. 4C). Another evidence of the importance of neighboring proteins to Tn5 cleavage is the fact that the relative predictive power of ATAC-seq in relation to DNase-seq varies for particular TF families. This includes TFs from the bZIP and helix-loop-helix families, which bind as dimmers and have large structures. These structural properties are likely to impair access of Tn5 to neighboring DNA regions.

HINT-ATAC is the only footprinting method exploring both nucleosome number decomposition and strand specific cleavage bias for improving footprinting prediction. Indeed, our comparative analysis indicate a clear advantage of HINT-ATAC in prediction of footprints in relation to state-of-art footprinting methods DeFCoM, Wellington, DNase2TF, PIQ and HINT-BC. The advantage of HINT-ATAC is also clear on libraries without paired-end sequences, where nucleosome number decomposition is not possible. Nucleosome decomposition but not strand specific signals were already used for nucleosome positioning in DNase-seq (Vierstra et al. 2014). While our analysis reinforces the importance of strand specific signals for all DNase-seq protocols, we could not evaluate the impact of nucleosome architecture decomposition given the lack of paired end DNase-seq libraries for cells with several TF ChIP-seq data, as the ones evaluated in this study.

Altogether, footprints predicted in Omni-ATAC obtained better performance than standard and fast-ATAC protocols. As seen in Fig.3A, Omni-ATAC libraries have higher amount of fragments on peaks associated to mono and di-nucleosomes, while it has less fragments between the peaks of **`Nfr`** and **`1N`** or with large size (>500). This suggests that improvements introduced in Omni-ATAC protocol enrich for mono and bi nucleosome reads, which leaves more attenuated strand bias profiles in 1N and +2N reads than standard ATAC-seq (Fig. 4). Indeed our model selection experiments confirm that using separated signals for nucleosome free and nucleosome containing fragments is the best strategy for finding footprints in Omni-ATAC, while only nucleosome free reads are best for standard ATAC. Finally, we show that the footprint predictive performance is equivalent between Omni-ATAC-seq and DNase-seq protocols, despite some TF specific differences. This indicates that improvements in ATAC-seq protocols and computational approaches make it a competitive alternative to DNase-seq for identifying TF binding sites, even for experiments based on moderate number of reads (~50 millions) and low number of cells (~50.000).

## Acknowledgements

We would like to thank Eduardo Gadde Gusmao (Harvard Medical School), Alvaro Rada-Iglesias (University of Cologne) for discussions; and Jens Keilwagen (Julius Kühn-Institut) for support with the Slim model code. This work has been funded by the Interdisciplinary Center for Clinical Research RWTH Aachen Medical Faculty (IZKF Aachen) and by the Deutsche Forschungsgemeinschaft (DFG-GE 2811/3). Simulations were performed with computing resources granted by ITC RWTH Aachen University under project rwth0233.

## Footnotes

1 Query performed using the words “ATAC-seq” and “DNase-seq” at 11/16/2017 considering the number of series and samples deposited within the last year

